# Two-Photon Microperimetry: A media opacity-independent retinal function assay

**DOI:** 10.1101/2020.06.27.175315

**Authors:** Ang Wei, Urmi V. Mehta, Grazyna Palczewska, Anton M. Palma, Vincent M. Hussey, Luke E. Hoffmann, Anna Diep, Kevin Nguyen, Bryan Le, Steven Yone-Shun Chang, Andrew W. Browne

## Abstract

Humans perceive light in the visible portion of electromagnetic radiation. However, visible light is scattered and attenuated by optical media opacities. Because all conventional visual function tests rely on visible light, test results are reduced in patients with optical media opacities like corneal scars, cataracts, and vitreous hemorrhages. Infrared (IR) light has greater penetrance through tissue than visible light. Two-photon IR visual stimulation, a recently pioneered technology, should enable testing of retinal visual function and produce results that are less susceptible to media opacities. The effects of simulated media opacities on visual performance in young healthy volunteers and the change in visual function in healthy phakic patients of two age ranges (20-40 and 60-80-year-old) were studied using conventional testing and 2-photon infrared visual stimulation. All subjects completed visual function testing using cone contrast threshold (**CCT**) testing, conventional microperimetr*y*, visible light microperimetry from a novel device (**2PM-Vis**), and infrared 2-photon microperimetry (**2PM-IR**). Retinal sensitivity measured by 2PM-IR demonstrated lower variability than all other devices relying on visible spectrum stimuli. Retinal sensitivity decreased proportionally with the transmittance of light through each filter. CCT scores and retinal sensitivity decreased with age in all testing modalities. Visible spectrum testing modalities demonstrated larger test result differences between young and old patient cohorts; this difference was inversely proportional to the wavelength of the visual function test. 2PM-IR mitigates media opacities which may mask small differences in retinal sensitivity when tested with conventional visual function testing devices.

**One Sentence Summary:** Two photon infrared visual function testing produces results that are less susceptible to media opacities than conventional tests.

## Introduction

Visual function is fundamental to eye care. Accurate measurements can identify visual disorders prompting medical intervention to preserve and restore vision. The highest acuity and color vision is subserved by the fovea located centrally in the light sensitive retina. The fovea is densely populated with 3 types of cones that are responsible for central visual function. Cone function is evaluated subjectively with visual acuity testing(*1, 2*), color and contrast testing(*3, 4*), and microperimetry(*5*). All conventional tests of visual function use visible light and their results are altered by ocular media opacities such as cataracts, corneal scars, and dense vitreous debris. A visual function test that is less susceptible to media opacity would produce more reliable information about retinal function in common blinding diseases. This would be most useful in diseases with concomitant cataract like age-related macular degeneration, myopic degeneration, diabetic retinopathy, and glaucoma.

Visual acuity (**VA**) testing is the most common method for evaluating central visual function and is the “gold standard” for assessing functional primary outcomes in clinical trials(*1*). VA testing is often performed using the Early Treatment Diabetic Retinopathy Study (**ETDRS**) or Snellen charts. However, VA is a poor predictor of total visual function(*6*), especially in patients with macular disease, as it provides no information on contrast sensitivity, color vision, nor metamorphopsia. Additional information for central visual function is available with cone contrast threshold (**CCT**) testing and microperimetry. CCT is an adaptive computer-based program that quantifies the severity and type of color vision deficiency (**CVD**)(*3, 4*) by selectively stimulating L-cones, M-cones, and S-cones. Microperimetry is a retinal sensitivity assay that provides quantitative retinal sensitivity data for both central and peripheral microscopic foci in the retina(*5*). This data is superimposable on real-time fundus images to correlate abnormalities in retinal structure with function.

These conventional visual function tests rely upon visible spectrum stimuli, typically described as ranging from 400 to 700 nm, and like all subjective visual function tests, suffer diminished reliability in patients with media opacities. VA testing using ETDRS charts has proven to be a poor predictor of macular function in patients with macular edema secondary to retinal vascular disease, in part due to anterior or posterior segment opacities(*7*). Cataractous lenses decrease the sensitivity threshold of all three cone classes due to increased scatter and decreased transmittance of visible light(*8–11*), with the greatest impact on short-wavelength cones especially after the age of 60(*11, 12*). Because of this effect, Fujikawa et al. established that CCT can be used reliably up to the seventh decade of life and after cataract surgery in elderly patients(*13*). Cataractous lenses and posterior capsular opacification (*14*) also diminish retinal sensitivity measurements obtained using microperimetry. This decrease is dependent upon the density and type of cataract(*6*) formed.

Boettner and Wolter showed that scattering decreases and transmittance increases as the wavelength elongates from visible to infrared (**IR**) ranges(*15*). Therefore, if humans perceived IR light, visual function testing would be less affected by cataracts and other media opacities. Previously, a 2-photon (**2P**) microperimeter demonstrated that IR laser light can initiate phototransduction when visual pigments absorb 2-photon light, inducing photoisomerization of 11-cis-retinal (opsins in rod and cone photoreceptors)(*16, 17*). Therefore, 2P IR stimulation of the retina provides an opportunity to evaluate retinal function and quantify visual function in a manner less susceptible to media opacity.

To explore this topic, a two-photon microperimeter (**2PM**) using pulsed IR light (**2PM-IR**) at 1045 nm was compared with United States Food and Drug Administration (US FDA) approved clinical visual function devices: CCT and conventional microperimetry (cMP). The 2PM device can also produce a visible 522.5 nm stimulus (**2PM-Vis**), which was also compared with 2PM-IR. Young healthy volunteers performed testing on all 4 devices using different optical filters that were placed directly in front of their eyes. A second analysis was performed in two separate cohorts comparing the effect of age on test performance using conventional and novel visual function tests. 2PM-IR visual stimulation was hypothesized to produce visual sensitivity results that are more consistent when tested in healthy subjects of all ages with and without filters than the 3 visible light visual function assays: CCT, cMP, and 2PM-Vis.

## Results

### Filter spectral transmission and CCT screen emission

Figure 1 displays the spectral emission of the CCT monitor as color-coded shaded histograms, transmission spectrum for media opacity-simulating filters as line graphs, and peak opsin spectral absorbances as horizontal bar plots. The spectral emission ranges for blue, green, and red light are: 418 nm-556 nm (peak 445 nm), 475 nm-615 nm (peak 541 nm), and 577 nm-724 nm (peak 608 nm), respectively. The peak spectral absorbances for rods, L-cones, M-cones, and L- cones are: 498 nm, 420 nm, 534 nm and 564 nm respectively(*18*).

**Figure 1.**
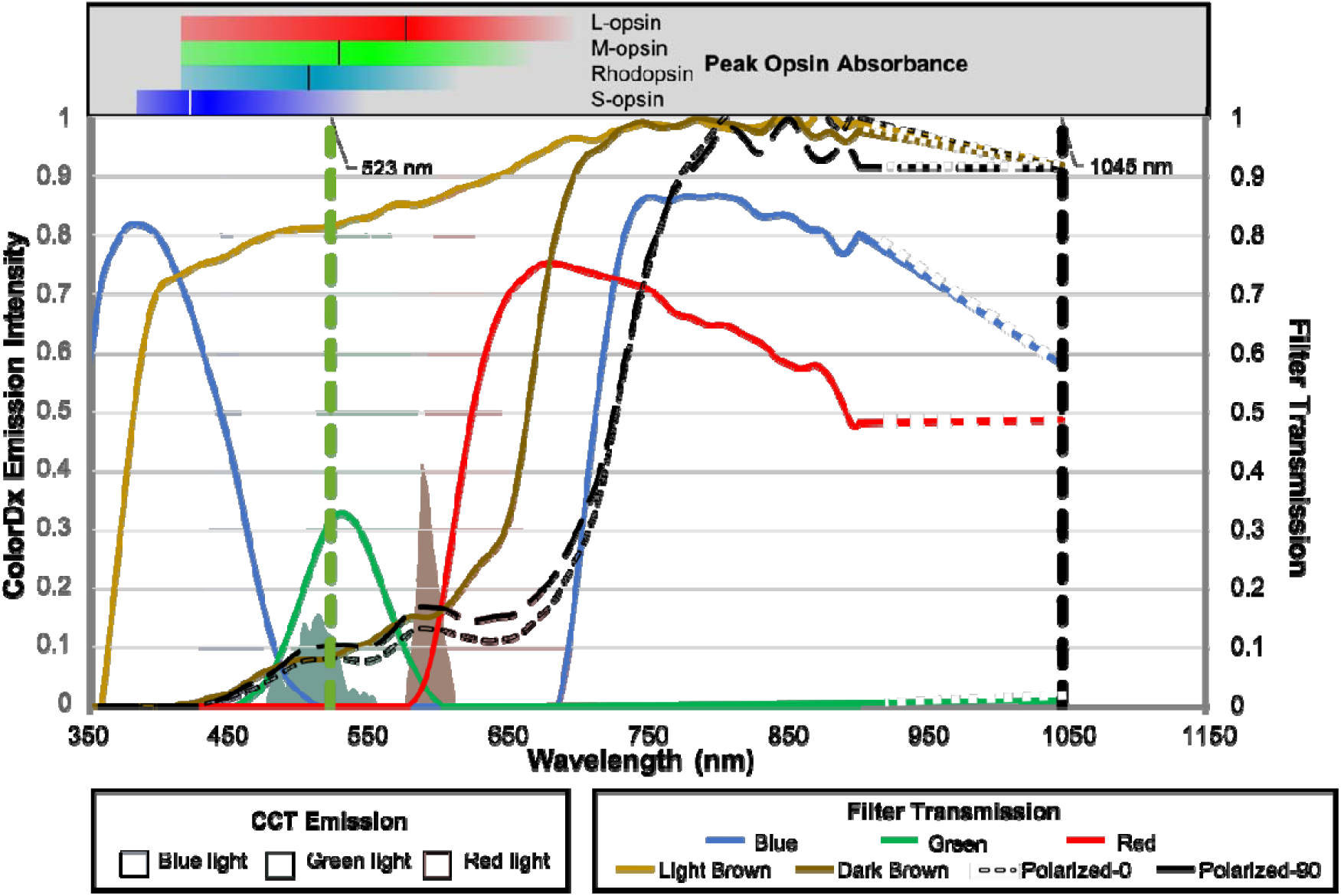
Spectral transmission of filters (lines) superimposed on the spectral emission of RGB light from the CCT monitor (shaded regions). Horizontal bars above the plot represent opsin spectral absorbances with vertical lines highlighting peak spectral absorption(*18*). Vertical dashed lines indicate wavelengths of visible green 2PM-Vis (523 nm) and pulsed infrared 2PM-IR (1045nm) stimulation.

The green filter transmits the least light across the full spectrum with low transmission even in the green spectrum centered around 531 nm and, notably, no transmission of IR light. The blue filter has maximal transmission in the blue spectrum centered around 379 nm and permits transmission of some IR light. The red filter has maximal transmission in the red spectrum centered around 681 nm and permits the transmission of IR light. The light brown filter minimally attenuates shorter wavelengths in the UV-blue spectrum, while the dark brown and black polarized filters significantly attenuate the entire visible spectrum. The light brown, dark brown, and dark black polarized filters all permit IR transmission at 1045nm.

### Microperimetry

Retinal sensitivity was greatest when no filter was used for all three perimetry tests (**Fig. 2**). Each filter condition decreased retinal sensitivity to a degree dependent on its transmission with p-values less than 0.001 for 2PM-IR and 0.018 for 2PM-Vis (**Fig. 3**). 2PM-IR sensitivity was most attenuated by the green filter, while marginally reduced by the blue and red filters. Light brown, dark brown, and black polarized filters marginally reduced retinal sensitivity. 2PM-Vis sensitivity was greatest with the green and light brown filters and lowest with the red filter. Blue and red filters severely diminished retinal sensitivity results on cMP. The green filter blocks IR light, preventing cMP from registering eye movement. Therefore, no data was recorded for this filter.

**Figure 2.**
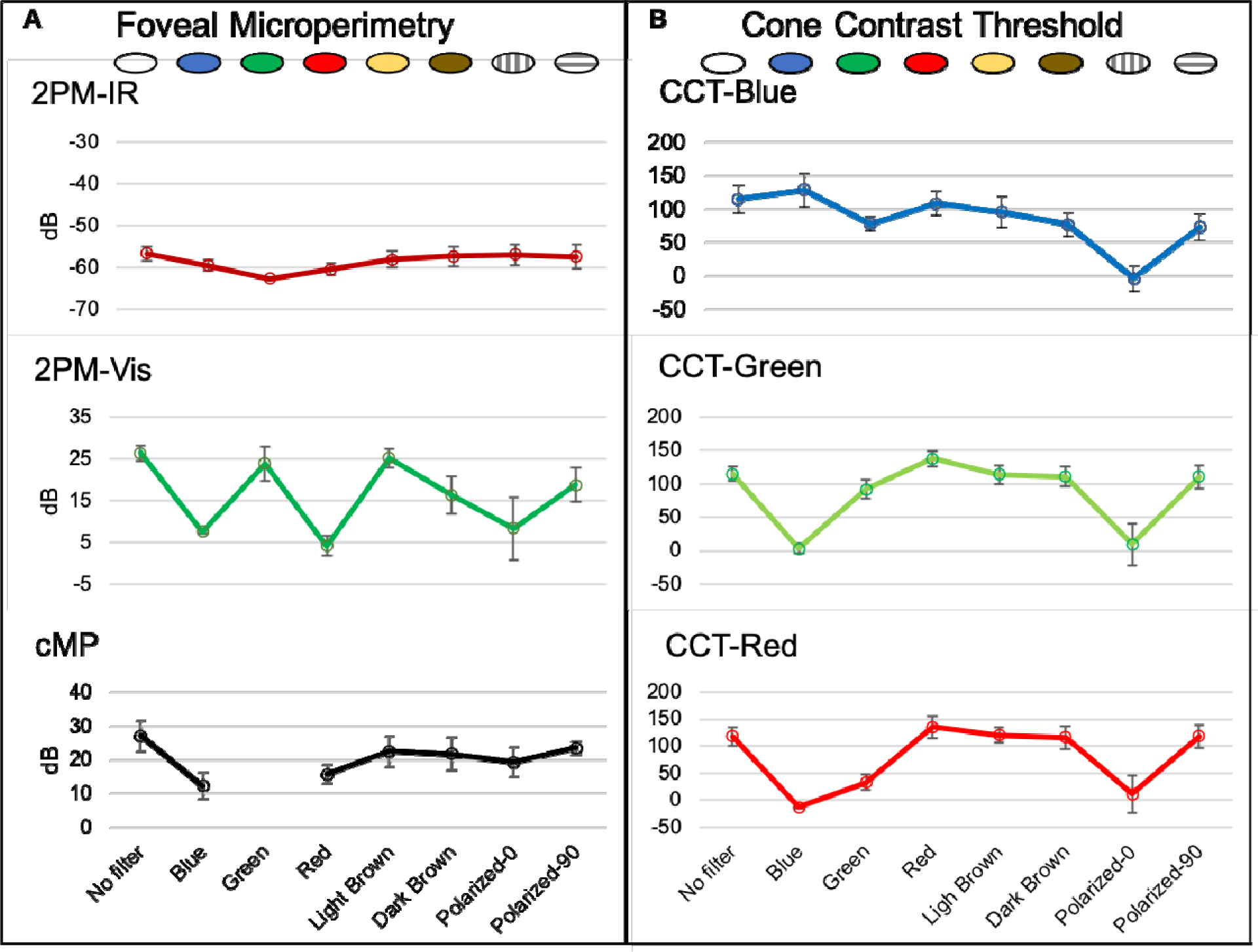
Effect of filters on visual function testing: (**A**) Foveal microperimetry with 2PM-IR, 2PM-Vis, and cMP testing. (**B**) Cone contrast threshold testing for each opsin.

**Figure 3.**
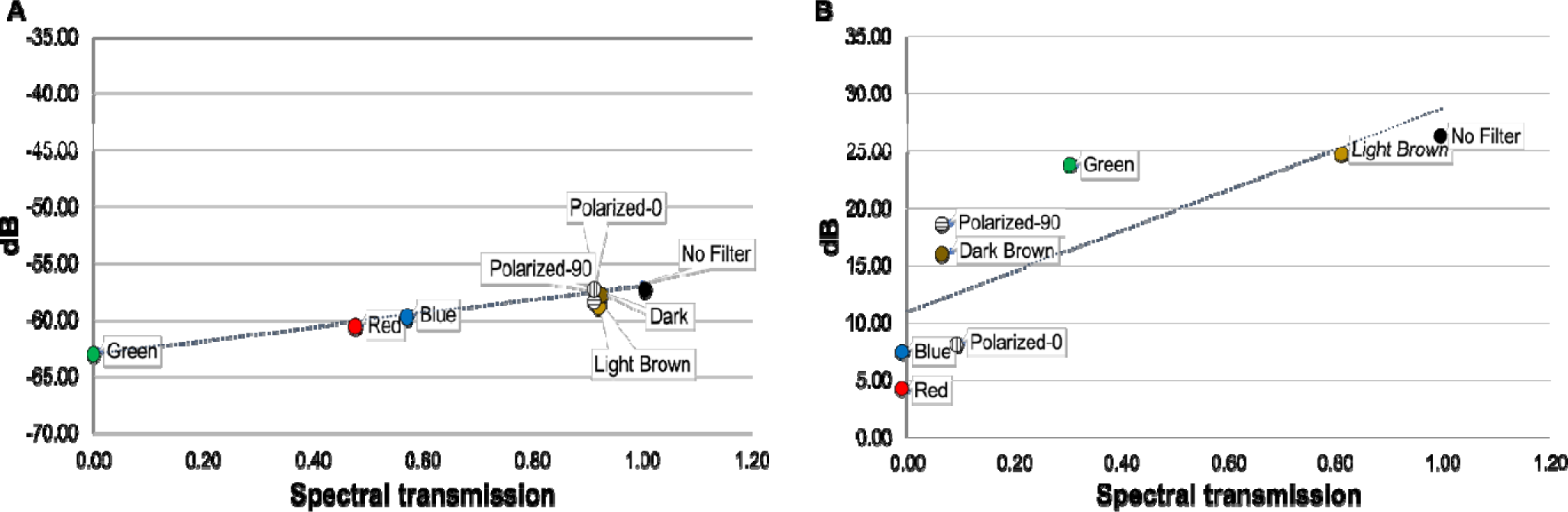
Linear regression of averaged retinal sensitivities versus filter spectral transmission for (**A**) 2PM-IR (R_2_= 0.965, β=−62.967, P<0.001) and (**B**) 2PM-Vis (R_2_= 0.637, β=11.002, P=0.005).

### Cone Contrast Threshold

All 3 cones were most sensitive when their respective color filter (RGB) was applied (red filter to stimulate L-opsin, green filter to stimulate M-opsin, and blue filter to stimulate S-opsin). The shaded filters (light brown, dark brown and dark black polarized filters) maximally reduced blue cone sensitivity, with only a nominal reduction in red and green cone sensitivities. The black polarized filter had the greatest reduction in all cone classes when oriented 0 degrees parallel to the polarization of the CCT liquid crystal display.

### Variability of perimetry and CCT measures

Although CCT is not a perimetry assay and reports sensitivity as a unitless Rabin performance score*(3)*, variability in results were similar to that seen for visible light perimetry (2PM-Vis and cMP). The sensitivity range for cMP, 2PM-Vis, and 2PM-IR were 11.33 dB (SD 4.96), 22.17 dB (SD 2.14), and 5.99 dB (SD 0.87) respectively.

### Mydriatic vs non-mydriatic

Table 1 reports the effect of mydriasis on retinal sensitivity. Test subjects performed 2PM-Vis and 2PM-IR testing in the non-mydriatic and mydriatic states. Mydriasis had a nominally positive but statistically insignificant effect on retinal sensitivity.

**Table 1.** Effect of mydriasis using 2PM. Paired t-tests were performed to assess the effect of mydriasis on retinal sensitivity. Data is represented as mean (SD) unless otherwise noted.

|  | No Mydriasis (dB) | Mydriasis (dB) | Mean difference (95% CI) | P-value |
| --- | --- | --- | --- | --- |
| **IR** | -57.89 (2.31) | -56.50 (3.33) | 1.39 (-0.09, 2.87) | 0.06 |
| **Vis** | 24.31 (5.67) | 26.45 (2.03) | 2.14 (-2.56, 6.83) | 0.30 |

### Effects of age on test performance

Results from visual function testing in young and old subjects are reported in figure 4. Nineteen subjects (33 eyes) between the ages of 20-40 were included in the “young” group and 23 subjects (32 eyes) between the ages of 60-80 were included in the “old” group. The distributions of visual function test scores by age group are displayed using violin plots. The effect of age is represented by the means and 95% confidence intervals for each group (grey lines and bars, respectively). CCT scores and retinal sensitivity were lower in older subjects across all testing devices. The rate of decline between young and old groups was greatest for S-cones: slope for each year increase in age −1.49 (95% CI: −1.9, −1.2) on CCT and least for 2PM-IR: slope −0.6 (95% CI: −1.4, −0.5). M- and S-cone function differences between the two age groups were sequentially smaller than the S-cone class and were inversely proportional to wavelength. Differences between young and old groups are statistically significant across all testing modalities. The difference was greatest for the S-cone CCT and least for 2PM-IR. Violin plot heights are shorter and more compact for both age groups when assessed by 2PM-IR than 2PM-Vis and CCT.

**Figure 4.**
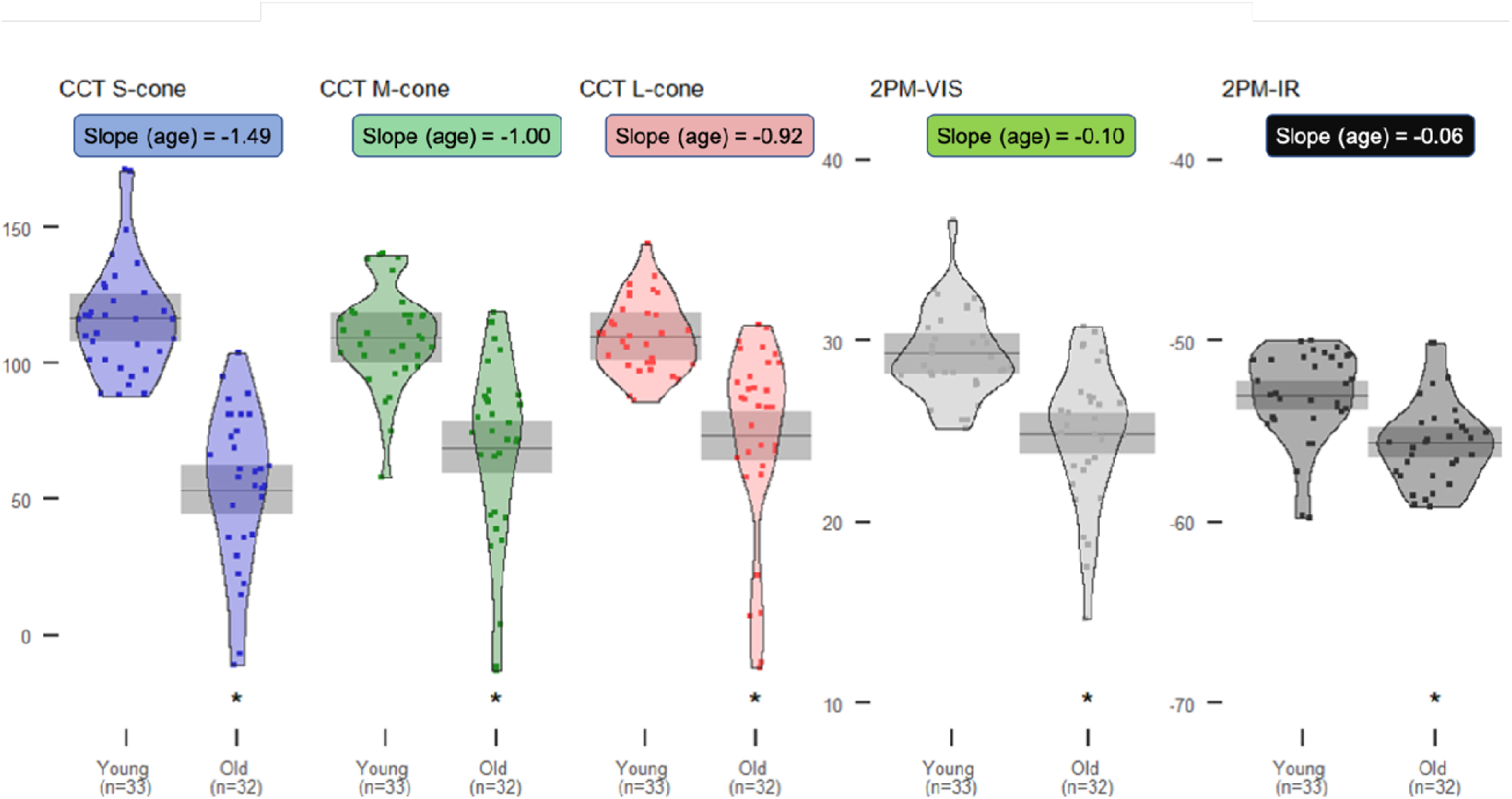
Color vision score distributions and means (95% CI) by age. Cone contrast threshold (CCT) scores and retinal sensitivity were assessed using conventional and novel visual function testing devices between young (20-40 y/o) and old (60-80 y/o) healthy subjects. Asterisks indicate statistically significant differences (p < 0.05) between younger and older groups. Slope indicates decrease in score per year increase in age for each test, modeled using linear regression.

## Discussion

Test performance comparing 2 US FDA-approved visual function tests (CCT and cMP) and 2 novel microperimetry assays (2PM-IR and 2PM-Vis) were evaluated in 2 separate studies. Young healthy subjects performed testing under 9 unique optical conditions. CCT scores and retinal sensitivity were negatively affected by all 7 filters. However, results obtained using 2PM-IR produced results with the least variability (Fig. 2 and 4). 2PM-IR regression plots were nearly flat and independent of visible spectrum optical transmission (Fig. 3), indicating a near independence from opacities, like cataracts, which attenuate light in the visible spectrum. Results obtained from CCT were highly variable and dependent on media filters, highlighting that it may not be a reliable assessment of macular health in patients with media opacity as was previously noted (*13*).

Retinal sensitivity assessed via 2PM-IR and cMP was proportional to the transmission of IR light through each filter (Fig. 2a). The green filter completely blocked transmission of IR light, thus preventing cMP from registering retinal landmarks and diminished 2PM-IR retinal sensitivity (Fig 1, 2a and 3a). Retinal sensitivity using 2PM-Vis was proportional to the transmission of green light through each filter (Fig 1, 2a and 3b). The red filter permitted negligible transmission of 522.5nm light and subjects were unable to detect the green stimulus pattern. As anticipated, the light brown filter had the highest transmission rate at 522.5 nm and produced results with the highest retinal sensitivity.

CCT values were proportional to the amount of overlap between the LCD display’s spectral emission and transmission of each filter (Fig. 1). The red filter overlaps the spectral emission of both the red and green LCD pixels and color sensitivity was concordant for the red and green cones. The green filter produced greatest color sensitivity for the green and blue cones as both these cones have overlapping absorbance curves that overlap with the green filter spectral transmission.

CCT scores and retinal sensitivity declined with age across all visual function testing devices in phakic subjects. However, the difference in score reports between young and old subjects decreased as the wavelength of the stimulus projected onto the retina increased, indicating that light in the near-IR spectrum has greater transmittance and penetrance through the aged eye than shorter wavelength light. CCT results demonstrated the expected trend of longer wavelengths entering the eye with less attenuation as S-cones showed the greatest difference between young and old subjects, while L-cones showed the smallest difference. Similarly, retinal sensitivity obtained using 2PM-Vis showed larger differences between the 2 groups compared to 2PM-IR, indicating that visual function testing using IR light is less affected by lenticular senescence and cataract formation. Furthermore, the violin plots for 2PM-IR are shorter for both age groups compared to 2PM-Vis, highlighting that IR light is less absorbed and less scattered than visible light, diminishing test variability(*17*). In this work, both 2PM-IR stimulation at 1045nm and single photon 2PM-Vis stimulation at 522.5 nm were perceived as a green stimulus. Because 2-photon infrared stimulation results in a perceived stimulus that is half the wavelength of the IR stimulus(*16*), 2PM-IR can be implemented to test the sensitivity of each opsin.

This is the first report comparing a novel visual sensitivity assay with established conventional visual function tests. IR light is absorbed less by ocular media and therefore penetrates deeper into the eye than visible spectrum light. Traditional visual function tests suffer diminished reliability in patients with lenticular senescence and media opacities(*6, 13, 14*). Ruminski et al previously demonstrated greater light scattering and attenuation for 2PM-Vis and less scattering and attenuation for 2PM-IR(*17*). Our study confirmed that a series of filters simulating various media opacities and normal aging diminish retinal sensitivity to a greater extent using visible spectrum tests compared to 2PM-IR. A clinical device employing 2PM-IR stimulation would reduce the impact of media opacity on overall visual function and better discern retinal and optic nerve performance.

A limitation in our study evaluating the effects of different transmission filters on retinal sensitivity is our small sample size and limited age range. However, this was intentional to demonstrate the impact of media opacity on visual function tests in young healthy eyes. Brown filters simulating media opacity are most clinically relevant because they mimic cataract. CCT scores were only significantly diminished for the S-cone class, and nominally reduced for M and L-cone classes and this recapitulates findings (Mehta et al. Under Review TVST). It should be noted, however, that CCT, which is very different from microperimetry, and cMP are not scotopic. Both CCT and cMP showed nominal reductions in sensitivity with brown filters. Conversely, 2PM is scotopic and the effect of brown filters was only significant for 2PM-Vis and not for 2PM-IR. This may suggest that the most sensitive and reliable visual function test is scotopic microperimetry using IR stimulation. Further investigation stratifying 2PM-IR values in different disease states are warranted. Additional ergonomic adjustments to the 2PM will improve ease-of-testing for the subject, especially in diseases causing poor fixation.

In conclusion, 2PM-IR can be implemented as a next-generation visual function platform that is nearly impervious to media opacity. This would have the greatest utility for studying diseases with a higher prevalence in the aging population, most notably glaucoma and age-related macular degeneration.

## Materials and Methods

This study received institutional review board approval from the University of California, Irvine and was conducted in accordance with the Declaration of Helsinki. All participants provided written informed consent before testing began. Two prospective studies evaluated the effects of various testing conditions and age on test performance.

### Subjects and Protocol

#### Effects of filters and mydriasis on visual function testing

The effects of various testing conditions on test performance were evaluated in this prospective study. Six phakic eyes of 6 healthy subjects (4 males, 2 females, age 23-29) underwent visual function testing using 4 different visual function devices: CCT (ColorDx CCT HD, Konan Medical, Irvine CA), conventional microperimetry (**MP-3**, Nidek Inc, San Jose, CA), visible light microperimetry produced by the novel 2PM device (**2PM-Vis**) and 2PM-IR. Figure 5a displays the novel 2PM apparatus for measuring retinal sensitivity. Nine separate tests were performed on each device under the following conditions: no filter, red filter, green filter, blue filter, light brown filter, dark brown filter, polarized black filter (0-degree rotation), and polarized black filter (90-degree rotation) (Fig. 5b). Subjects subsequently performed 2PM-IR and 2PM-Vis testing without a filter in the mydriatic state following instillation of 0.75% tropicamide and 2.5% phenylephrine.

**Figure 5.**
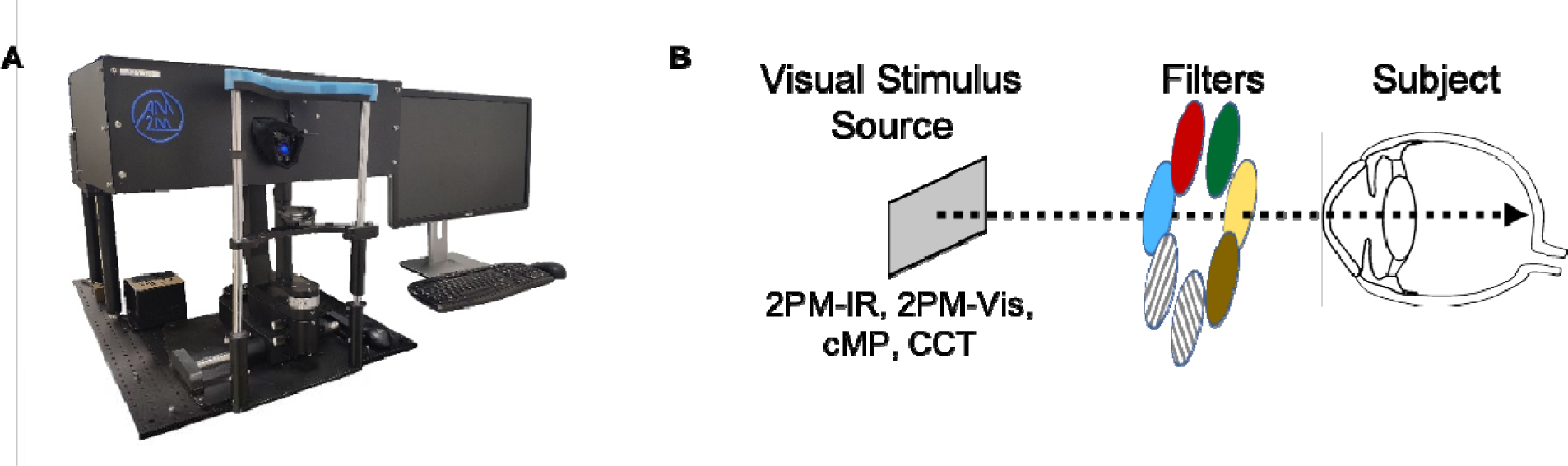
2PM is capable of testing retinal sensitivity using single photon visible (2PM-Vis) and 2-photon infrared (2PM-IR) stimuli **(A)**. Configuration for measuring cone contrast/retinal sensitivity under the 9 different testing conditions (**B**).

#### Effects of age on test performance

The effects of age on test visual function was evaluated in 42 healthy subjects. Nineteen subjects (33 eyes) between the ages of 20-40 and another 23 subjects (32 eyes) between the ages of 60-80 were analyzed using CCT, 2PM-Vis, and 2PM-IR.

Inclusion criteria for both analyses were: (1) age >18 years, (2) no history of ocular disease, (3) BCVA of 20/25 or better, and (4) phakic lens status. Normal ophthalmic health was determined by clinical examination by a retina specialist and review of color fundus photography (Clarus, Carl Zeiss Meditec, Dublin, CA) and spectral domain optical coherence tomography (Cirrus 5000, Carl Zeiss Meditec) demonstrating normal anatomy. Visual acuity was measured following ETDRS testing protocol.

#### Spectral Emission and Transmission

The spectral transmission of each filter was measured using a compact spectrometer (CCS200/M, Thorlabs, Newton, US) in indirect sunlight and quantified from 350-900 nm. Filter transmission at 1045nm was quantified using a 1045 nm laser source. The spectral emission of RGB light from the ColorDx liquid crystal display (LCD) was measured using the same compact spectrometer. The spectral emission from cMP was not obtained due to difficulty probing light in the device oculars but is assumed to be similar to the ColorDx LCD.

### Functional Testing

#### 2 Photon Microperimetry

2PM (Fig 5a, Polgenix, Cleveland, OH) is a custom apparatus that is the product of the National Eye Institute’s Audacious Goal Initiative to develop new technology for the non-invasive assessment of retinal structure and function in vivo. 2PM is a retinal sensitivity assay capable of stimulating the retina using pulsed 2-photon infrared (IR) light at 1045 nm (2PM-IR) or standard single-photon visible green light at 522.5 nm (2PM-Vis) to evaluate foveal sensitivity. Testing was performed in a dimly lit room using the subject’s BCVA after 30 minutes of dark adaptation. A stimulus pattern formed either by 2P infrared or visible light was projected onto the retina as previously described(*17*). Briefly, a pulsed 2P IR signal was generated as a 63 MHz pulse train of 250 fs-long pulses with a 1045 nm central wavelength. Visible (green) light at 522.5 nm was produced by diverting the IR light source to a non-linear crystal producing a half wavelength change of the 1045nm laser source to a 522.5 nm visible stimulus(*17*). Test subjects were asked to adjust the intensity of the stimulus by either scrolling up or down on a computer mouse, which would either increase or decrease the intensity of the stimulus entering the eye. Once the visibility threshold was reached and the subject could no longer see the stimulus, they were asked to left click on the mouse, thereby recording the threshold power that could be detected. The same procedure was performed 4 more times and an average threshold sensitivity for all 5 replicate tests was calculated. Small refractive errors were corrected using correction lenses internal to the 2PM.

2PM-IR and 2PM-Vis stimulus intensity are measured in microwatts and femtowatts respectively. Retinal sensitivity (S) was calculated using the inverse of the average threshold power (S= 1/threshold power). This value (P_1_) was then converted to decibels using 100 pW (P_0_) of threshold power as the reference value using the following equation(*17*):

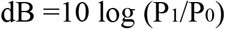

#### Conventional Visible Microperimetry

Conventional microperimetry was performed using the MP-3, a clinical microperimetry assay used to quantify macular sensitivity. It was performed only for the foveal point under photopic conditions without correction. The contralateral eye was occluded. Goldmann III stimuli were applied over an intensity range from 0-34 dB. An IR sensor was used to track eye movement and a white LED light source was used to project a stimulus pattern onto the retina.

#### Cone Contrast Threshold Testing

The ColorDx CCT HD is an adaptive visual function testing device that selectively stimulates retinal L, M, and S-cones. A series of Landolt C optotypes are projected in either decreasing or increasing steps of cone contrast against an isoluminant grey background(*19*). Subjects indicate the orientation of the gap in the Landolt C optotype using arrows on a trackpad. The final score was calculated based on the number of correct answers. CCT was performed monocularly using the subject’s BCVA under photopic conditions at a distance of 2 feet. Score reports are as follows: >90 is normal, 75-90 indicates possible color vision deficiency (CVD), <75 indicates CVD.

#### Statistical Analysis

Mean sensitivities and standard deviation were calculated for each test. Paired sample t-tests were performed to assess whether mydriasis affected retinal sensitivity.

Linear regression models were fit to compare the effect of age between young and old subjects on visual function test performance assessed using CCT, cMP, 2PM-Vis, and 2PM-IR. Standardized coefficients and 95% CIs were used to estimate the effect size between young and old age groups for each testing modality since unit scales varied between devices. Standard coefficients can be interpreted as the differences in standard deviations between young and old groups. Test score distributions by age group were illustrated using violin plots. All statistical analyses were performed using IBM SPSS Statistics 25 and R version 3.6.1.

## Acknowledgments

The authors would like to thank Dr. Grazyna Palczewska and Dr. Krzysztof Palscewski for use of the 2PM. The authors would also like to thank Carl Zeiss Meditec for loan of ophthalmic testing devices used to verify ophthalmic health in human subjects and Konan medical for an unrestricted donation of ColorDX.

## Author contributions

All authors played a role in writing and revising the manuscript. A.W, U.V.M, V.M.H, L.E.H, A.D, K.N, B.L, and S.Y.C conducted the experiments. A.M.P. performed the statistical analysis. A.W.B designed the experiments.

## Data and materials availability

All data related to this paper will be deposited in an approved database as excel files.

## References and Notes

1. Kaiser, P.K., Prospective evaluation of visual acuity assessment: a comparison of snellen versus ETDRS charts in clinical practice (An AOS Thesis). Trans Am Ophthalmol Soc, 2009. 107: p. 311–24.

2. Ahmed, S.F., K.C. McDermott, W.K. Burge, I.I.K. Ahmed, D.K. Varma, Y.J. Liao, A.S. Crandall, and S.K.R. Khaderi, Visual function, digital behavior and the vision performance index. Clin Ophthalmol, 2018. 12: p. 2553–2561.

3. Rabin, J., J. Gooch, and D. Ivan, Rapid quantification of color vision: the cone contrast test. Invest Ophthalmol Vis Sci, 2011. 52(2): p. 816–20.

4. Rabin, J., Quantification of color vision with cone contrast sensitivity. Vis Neurosci, 2004. 21(3): p. 483–5.

5. Laishram, M., K. Srikanth, A.R. Rajalakshmi, S. Nagarajan, and G. Ezhumalai, Microperimetry - A New Tool for Assessing Retinal Sensitivity in Macular Diseases. J Clin Diagn Res, 2017. 11(7): p. NC08–NC11.

6. Richter-Mueksch, S., S. Sacu, B. Weingessel, V.P. Vecsei-Marlovits, and U. Schmidt-Erfurth, The influence of cortical, nuclear, subcortical posterior, and mixed cataract on the results of microperimetry. Eye (Lond), 2011. 25(10): p. 1317–21.

7. Hatef, E., M. Hanout, A. Moradi, E. Colantuoni, M. Bittencourt, H. Liu, Y.J. Sepah, M. Ibrahim, D.V. Do, D.L. Guyton, and Q.D. Nguyen, Longitudinal comparison of visual acuity as measured by the ETDRS chart and by the potential acuity meter in eyes with macular edema, and its relationship with retinal thickness and sensitivity. Eye, 2014. 28(10): p. 1239–1245.

8. Nguyen-Tri, D., O. Overbury, and J. Faubert, The role of lenticular senescence in age-related color vision changes. Invest Ophthalmol Vis Sci, 2003. 44(8): p. 3698–704.

9. Paramei, G.V. and B. Oakley, Variation of color discrimination across the life span. Journal of the Optical Society of America a-Optics Image Science and Vision, 2014. 31(4): p. A375–A384.

10. Fristrom, B. and B.L. Lundh, Colour contrast sensitivity in cataract and pseudophakia. Acta Ophthalmol Scand, 2000. 78(5): p. 506–11.

11. Artigas, J.M., A. Felipe, A. Navea, A. Fandino, and C. Artigas, Spectral transmission of the human crystalline lens in adult and elderly persons: color and total transmission of visible light. Invest Ophthalmol Vis Sci, 2012. 53(7): p. 4076–84.

12. Ao, M., X. Li, W. Qiu, Z. Hou, J. Su, and W. Wang, The impact of age-related cataracts on colour perception, postoperative recovery and related spectra derived from test of hue perception. BMC Ophthalmol, 2019. 19(1): p. 56.

13. Fujikawa, M., S. Muraki, Y. Niwa, and M. Ohji, Evaluation of clinical validity of the Rabin cone contrast test in normal phakic or pseudophakic eyes and severely dichromatic eyes. Acta Ophthalmol, 2018. 96(2): p. e164–e167.

14. Varga, A., S. Sacu, P.V. Vecsei-Marlovits, S. Richter-Mueksch, T. Neumayer, B. Weingessel, O. Findl, and U. Schmidt-Erfurth, Effect of posterior capsule opacification on macular sensitivity. J Cataract Refract Surg, 2008. 34(1): p. 52–6.

15. Boettner, E.A. and J.R. Wolter, Transmission of the Ocular Media. Investigative Ophthalmology, 1962. 1(6): p. 776–783.

16. Palczewska, G., F. Vinberg, P. Stremplewski, M.P. Bircher, D. Salom, K. Komar, J.Y. Zhang, M. Cascella, M. Wojtkowski, V.J. Kefalov, and K. Palczewski, Human infrared vision is triggered by two-photon chromophore isomerization. Proceedings of the National Academy of Sciences of the United States of America, 2014. 111(50): p. E5445–E5454.

17. Ruminski, D., G. Palczewska, M. Nowakowski, A. Zielinska, V.J. Kefalov, K. Komar, K. Palczewski, and M. Wojtkowski, Two-photon microperimetry: sensitivity of human photoreceptors to infrared light. Biomed Opt Express, 2019. 10(9): p. 4551–4567.

18. Bowmaker, J.K. and H.J.A. Dartnall, Visual Pigments of Rods and Cones in a Human Retina. Journal of Physiology-London, 1980. 298(Jan): p. 501–511.

19. Medical, K. ColorDx CCT-HD Fundamentals. [cited 2020 March 1]; Available from: https://konanmedical.com/colordx-ccthd-fundamentals/.

